# Repli-seq: genome-wide analysis of replication timing by next-generation sequencing

**DOI:** 10.1101/104653

**Authors:** Claire Marchal, Takayo Sasaki, Daniel Vera, Korey Wilson, Jiao Sima, Juan Carlos Rivera-Mulia, Claudia Trevilla García, Coralin Nogues, Ebtesam Nafie, David M. Gilbert

**Author notes:** Correspondence should be addressed to DMG. Both authors contributed equally to this work (Claire Marchal, Takayo Sasaki).

## Abstract

Cycling cells duplicate their DNA content during S phase, following a defined program called replication timing (RT). Early and late replicating regions differ in terms of mutation rates, transcriptional activity, chromatin marks and sub-nuclear position. Moreover, RT is regulated during development and is altered in disease. Exploring mechanisms linking RT to other cellular processes in normal and diseased cells will be facilitated by rapid and robust methods with which to measure RT genome wide. Here, we describe a rapid, robust and relatively inexpensive protocol to analyze genome-wide RT by next-generation sequencing (NGS). This protocol yields highly reproducible results across laboratories and platforms. We also provide computational pipelines for analysis, parsing phased genomes using single nucleotide polymorphisms (SNP) for analyzing RT allelic asynchrony, and for direct comparison to Repli-chip data obtained by analyzing nascent DNA by microarrays.

## INTRODUCTION

DNA replication occurs during S phase of the cell cycle. In human cells, this process lasts around 8 hours^1^. Different regions of the genome replicate at different times during S phase, following a defined replication timing (RT) program ^2,3,4,5,6,7,8^. RT is highly conserved between mouse and human ^4,5,6,9^. Interestingly, in mammalian cells, almost 50% of the genome switches RT upon cell differentiation ^2,3,4,5,6,8^. RT is linked to transcription, although the causal links between these two processes are not clearly understood ^10,11,5,12,8,13^. Moreover, RT is closely associated with 3D nuclear architecture ^6,9^ as measured by chromatin conformation capture methods such as Hi-C 14. Early and late replicated regions correlate with the A and B compartments identified by Hi-C analysis ^5,6,14,15^, while domains of coordinately regulated RT (replication domains; RDs) align with topologically associating domains (TADs) 15. Accurate methods to analyze RT are essential to explore the links between all these processes.

Genome-wide RT analysis methods are based on the quantification of replicated genomic regions at different times during S phase. Multiple techniques have been developed to assess RT. One of the major applications, repli-chip, uses BrdU pulse labeling of nascent DNA, Fluorescence Activated Cell Sorting (FACS) to separate cells into different times during S phase ^16,17^, and BrdU immunoprecipitation to isolate newly synthesized DNA at different time points of the S phase ^2,18^. This newly synthesized DNA is then quantified by microarray hybridization. Subsequently, Hansen et. al. sequenced the newly synthesized DNA produced from BrdU labeling and FACS sorting to coin the term Repli-seq 4. In either case, the number of S phase fractions can be varied in this protocol, from 2 fractions (early vs. late or E/L), which produces a simple ratio of enrichment in early vs. late S phase, to multiple fractions (to date up to 8) of S phase ^4,16,17,19^ Both 2 and 6 fraction Repli-chip vs. Repli-seq give highly similar profiles after smoothing and normalization 15. Taking multiple fractions of S phase to date has provided little in the way of increased resolution, due to the fact that the labeling times with BrdU necessary for effective BrdU-IP (>60 minutes) label several hundreds of kilobases of DNA. Multiple fractions can, however, give information on synchrony of replication across a population of cells during S phase; highly asynchronous replication would give BrdU incorporation across S phase with multiple fractions but would average out to be indistinguishable from middle replication in a 2 fraction ratio method. When large numbers of different cell types or experimental conditions are being compared, however, the 2-fraction protocol is considerably faster and more robust in both wet and computational processing, provides highly reliable data with well established quality control standards and can be easily integrated in next generation sequencing analysis pipelines.

We previously published a genome-wide RT analysis protocol based on microarray hybridization: repli-chip 18. Here we summarize our newly adapted repli-seq protocol. While microarray processing can be less expensive both in wet and computational costs, sequencing of newly synthesized DNA in repli-seq provides distinct advantages. Sequencing allows one to overcome species limitations with microarray availability, and provides superior sensitivity to allele-specific microarrays in cases of studies needing sequence specificity such as distinguishing single nucleotide polymorphism (SNPs) / quantitative trait loci ^20,21,22^ or to compare homologous chromosomes with phased genomes.

We describe here the complete optimized repli-seq experimental procedures and bioinformatic analysis pipeline developed and routinely used in the lab for rapid high confidence analysis of samples containing as few as two thousand S phase cells. This protocol generates data to study RT at the genome-wide level, in a sequence specific manner. We have successfully applied this protocol to human and mouse embryonic stem cells (ESC), ESC-derived, primary cells, and cell lines, as well as frozen viable banked tissue samples ^15,23^ (and unpublished work). Moreover, this protocol could be easily adaptable to study other species and measure allele-specific RT if a phased genome is available.

### Overview

The experimental parts of this protocol begin with cultured cells and end with sequencing of the samples and are composed of 9 steps. This is followed by the analysis part of the protocol, which starts from the fastq or fastq.gz files obtained after sequencing and generates normalized bedgraph files that can be used for further bioinformatics studies. An overview of the protocol is presented in Figure 1.

**Figure 1:**
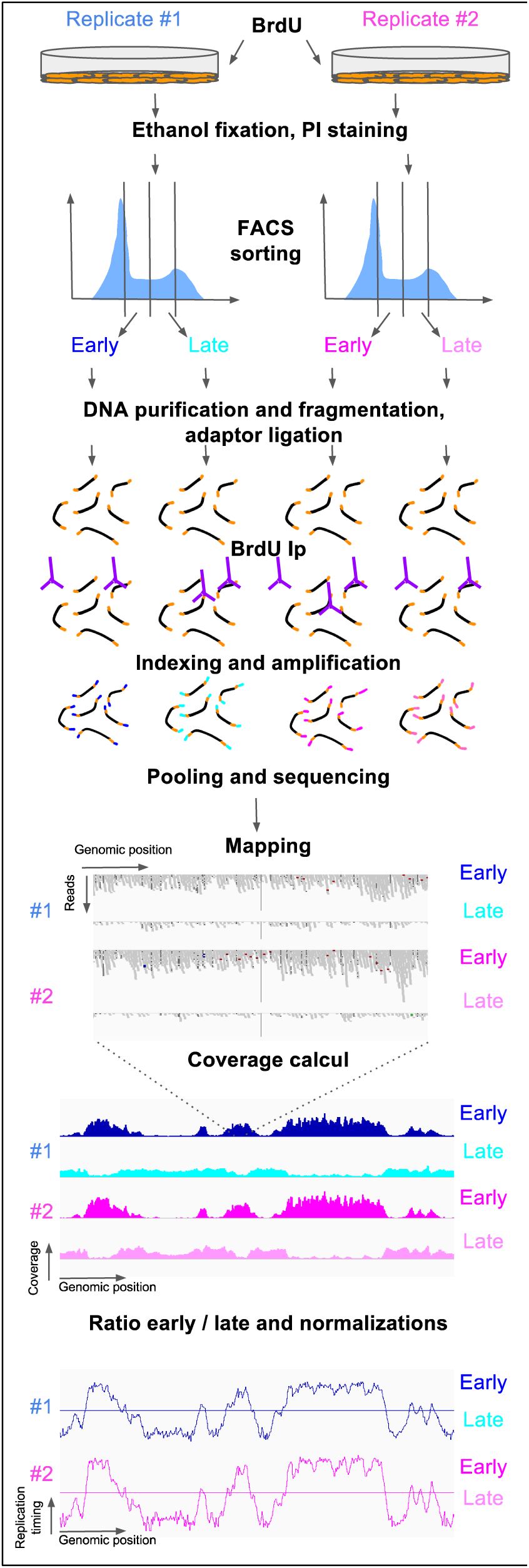
Overview of repli-seq protocol and analysis.

#### Labeling and fixation

This step is performed on cells that can be pulse-labeled, which includes any cell type that is proliferating and expresses a thymidine kinase, and that can be dissociated into single cells. Asynchronously proliferating cells are pulse-labeled with BrdU to mark newly synthesized DNA. The resolution of repli-seq is currently limited by the percentage of the genome that must be labeled for efficient BrdU immunoprecipitation (IP), which is approximately 10% of the genome. Since eukaryotic cells replicate their genomes as spatially clustered replicons, generally large domains are labeled in a short period of time 24. Improving IP signal to noise with shorter labeling periods (e.g. through the use of other labeling reagents) is an area of active exploration. If desired, BrdU can be replaced by EdU and the pull down performed by click chemistry, but to date we have not seen significant differences in the output using either label and BrdU-IP is less expensive and faster than EdU click chemistry. Labeling can be avoided entirely by using approaches that measure the <2 fold higher copy number of early vs. late replicating sequences, but the dynamic range of the data is reduced dramatically relative to nascent strand enrichment described here ^25,26^.

#### FACS sorting

BrdU-labeled cells are stained with propidium iodide (PI) to assess the cell cycle phase of each cell, through their DNA content. Cells are sorted by FACS according to their DNA content based on the PI staining, to isolate two cell populations: early S cells and late S cells. Many other DNA staining dyes may be substituted (e.g. chromomycin, mithramycin, DAPI, Hoechst) depending upon the fluorescence wavelengths and applications desired 27

#### DNA preparation and fragmentation

Genomic DNA from early and late S phase is purified and fragmented. DNA fragmentation is performed by sonication using a Covaris. The advantages of Covaris are: 1) it fragments DNA into a relatively tight size distribution reproducibly, hence there is no need for subsequent size selection most of the time; 2) the same conditions work for a broad range of DNA concentrations (50-5000ng). Although Covaris consumables are expensive, these advantages make the total cost of library construction lower than fragmenting DNA using other methods. The fragmentation is performed to obtain DNA fragments of an average length of 200bp if 50 to 100 bp single end sequencing is to be performed. Fragments size can be adapted for specific purposes. For example, for experiments where we wish to detect SNPs in phased genomes, we fragment to more than 500bp fragments and perform 250bp paired end sequencing.

#### Library construction

This step begins prior to the BrdU labeled DNA immunoprecipitation (BrdU IP). There are many advantages to constructing the libraries before the BrdU IP. First, the amount of available DNA is higher than after the IP and constructing libraries from small amounts of DNA is challenging. Second, BrdU IP generates single-strand DNA, which means that constructing the libraries after the BrdU IP would require converting it back to double-stranded DNA, which adds one more step to the protocol and could introduce artifacts. Importantly, this method also allows one to quantify the efficiency of BrdU IP by quantitative PCR of IP and input. The kit used to construct the library depends on the sequencer you will use. We use an NEB kit for Illumina sequencer. Libraries are constructed according to the manufacturer’s protocol, following three steps: end repair and dA tailing, adaptor ligation, and USER treatment.

#### BrdU Immunoprecipitation

DNA fragments, linked to the adaptors, are immunoprecipitated with an anti-BrdU antibody and a mouse secondary antibody.

#### Indexing and PCR amplification

DNA is indexed during the PCR amplification step, using an NEB kit. The optimum number of PCR cycles should be determined by qPCR if necessary.

#### Post-PCR Purification

DNA is purified to remove PCR reagents, primers and primer dimers, and proteins contamination. We use AMPure XP beads.

#### Quality Control, Pooling and Sequencing

Quality control is an important step to avoid sequencing low quality samples which could not be used for further analysis. This step includes the quantification of DNA concentration for each sample and analysis of the size distribution of the library. The performance of the BrdU-IP is assessed by quantitative PCR in known early and late replicating regions, if these data are available for your samples. After these quality control steps, libraries are pooled for sequencing. The pool of libraries is checked for size distribution and molar concentration before being sequenced. Sequencing is performed on a HiSeq illumina sequencer.

#### Analysis

Reads are mapped onto the genome using bowtie2. The coverage is assessed for each samples, and the base 2 log ratio of early vs. late S phase samples is calculated in genomic windows. All these steps can be performed using R software or in command line. Next, base 2 log ratio files are post-processed using R. Post-processing allows the comparison between samples when comparing samples with local RT changes. Post-processing includes quantile normalization and (optional) Loess smoothing. We do not recommend using quantile normalization when comparing datasets with global RT changes (such as Rif1 knockout 28). Loess smoothing can be skipped if you use overlapping genomic windows for coverage assessment. The analysis generates, for each sample, one bedgraph coverage file of post-processed log2 ratio of early vs. late S phase cells per sample. These files can be viewed using a genome viewer (IGV 29, IGB 30, USCS genome browser 31) or on the replication domain platform (www.replicationdomain.org), which allows comparison to our database of hundreds of available RT, genome-wide transcription and 4C datasets 32. Moreover, these files can be easily integrated into further analysis pipelines.

## MATERIALS

### REAGENTS

#### CRITICAL

All reagents/materials should be molecular biology / PCR grade.

- Cells of interest (see REAGENTS SETUP)
- BrdU (see REAGENTS SETUP)
- Cell culture medium and FBS appropriate for the cell type
- 1X Trypsin-EDTA (Mediatech 25-053-Cl) or other cell dissociation reagent appropriate for the cell type
- 1X PBS (Corning 21-031-CV)
- Propidium Iodide (PI) (Sigma P4170-) 1 mg/mL (see REAGENTS SETUP)
- RNase A 10mg/mL (Sigma R6513) Store at -20°C
- PBS / 1% FBS / PI / RNase A (see REAGENTS SETUP)
- Proteinase K 20mg/mL (Amresco E195) Store at -20°C
- SDS-PK buffer (see REAGENTS SETUP)
- 70% (vol/vol) Ethanol in H2O.
- Quick-DNA MicroPrep (Zymo D3021)
- 0.5 M EDTA pH 8 (Boehringer 808288)
- NEBNext Ultra DNA Library Prep Kit for Illumina (E7370) (see REAGENTS SETUP)
- TE (10 mM Tris pH 8.0, 1mM EDTA)
- 10 mM Tris pH 8.0 (Fisher BP152-5)
- 10X IP buffer (see REAGENTS SETUP)
- Anti-BrdU antibody (BD 555627) 12.5 μg/ml (see REAGENTS SETUP)
- Anti-mouse IgG ((Sigma M7023)
- Digestion buffer (see REAGENTS SETUP)
- NEBNext Multiple Oligos for Illumina (Dual Index Primers Set 1, Cat. # E7600S)
- DNA Clean & Concentrator-5 (Zymo Research D4014)
- Agencourt AMPure XP (Beckman Coulter A63880)
- Qubit^®^ dsDNA HS Assay Kit (Life Technologies Q32854)
- Agilent High Sensitivity DNA Kit (Agilent, 5067-4626)
- Agilent DNA 1000 Kit (Agilent, 5067-1504)
- 100% Ethanol (Sigma E7023)
- H_2_O
- Tween 20 (Sigma P1379) 0.05% (vol/vol) in 10 mM Tris pH 8
- KAPA Library Quantification kit (Kapa biosystems KK4824 for Applied Biosystems 7500 Fast)

#### EQUIPMENT

- Covaris E220
- microTUBE AFA Fiber Pre-Slit Snap Cap 6x16mm (Covaris 520045) (see EQUIPMENT SETUP)
- heat block
- Qubit
- Centrifuge (Eppendorf 5415D and Sorvall Legend RT)
- Magnet separator
- Thermal cycler (with plate for 0.5mL tubes)
- 1.5mL microcentrifuge tubes
- 0.5mL PCR tubes (we use Axygen PCR-05-C, but this reference has to be adapted to your thermal cycler)
- 0.65mL microcentrifuge tubes (Coster 3208)
- Real-Time Thermocycler (Applied Biosystems 7500 Fast)
- PCR plate (Life Technologies 4346906)
- Optical Adhesive Film (Life Technologies 4311971)
- Parafilm
- Vortex
- Unix-based Computer (see EQUIPMENT SETUP)

#### REAGENTS SETUP

- Cells of interest

Cultures can be grown in any size cell culture dish, but must be in an actively dividing state for use in this protocol. If you have to start from frozen samples that were proliferating at the time they were frozen but are no longer metabolically active, use the S/G1 method described by Ryba et al. 18. Since FACS can be problematic with a low cell number, we recommend starting with greater than 2 × 106 cells. We have successfully profiled RT using as low as 300,000 starting cells, and as few as 1000 early and late S phase cells after sorting, but cells are lost during PI staining, filtering, sorting so we do not recommend trying this without extensive experience.

CAUTION: All experiments should be performed in accordance with relevant health, safety and human subjects guidelines and regulations.

- BrdU (5-bromo-2’-deoxyuridine) (Sigma Aldrich, B5002) Make stock solutions of 10 mg/mL (and 1 mg/mL if you need to handle a small scale of culture) in ddH_2_O and store at -20°C in aliquots, protected from light.
- Propidium Iodide (1 mg/mL) (PI) To make 20 mL, dissolve 20 mg Propidium Iodide powder in autoclave ddH_2_O to achieve a final volume of 20 mL and filter. Store for up to one year at 4°C protected from light.
- PBS / 1% FBS / PI / RNase A Add 50 μl of 1 mg/ml PI, 25 μl of 10 mg/ml Rnase A to every 1 ml of PBS-1% FBS-SDS-PK buffer To make 50 mL, combine 34 mL autoclaved ddH_2_O, 2.5 mL 1M Tris-HCl pH 8.0, 1 mL 0.5 M EDTA, 10 mL 5 M NaCl and 2,5 mL 10% (wt/vol) SDS (Invitrogen 15525017) in H_2_O. Store at room temperature. Warm to 56°C before use to completely dissolve SDS.
- 10X IP buffer To make 50 mL, combine 28.5 mL ddH_2_O, 5 mL 1M Sodium Phosphate pH 7.0, 14 mL 5 M NaCl, and 2.5 mL 10% (wt/vol) Triton X-100 in H_2_O . Store at room temperature.
- anti-BrdU antibody 12.5 μg/ml Dilute antibody in 1X PBS from the stock concentration of 0.5 mg/mL to a final concentration of 12.5μ g/mL. Prepare 40μ l of diluted antibody for each sample and discard unused diluted antibody.
- Digestion buffer To make 50 mL, combine 44 mL autoclaved ddH_2_O, 2.5 mL 1M Tris-HCl pH 8.0, 1 mL 0.5 MEDTA, and 2.5 mL 10% (wt/vol) SDS in H_2_O. Store at room temperature.
- NEBNext Ultra DNA Library Prep Kit for Illumina (E7370) The Ligation Master Mix and Ligation Enhancer can be mixed ahead of time and the mix is stable for at least 8 hours at 4°C. We do not recommend premixing the Ligation Master Mix, Ligation Enhancer and adaptor prior to use in the Adaptor Ligation Step.

#### EQUIPMENT SETUP

- microTUBE AFA Fiber Pre-Slit Snap Cap 6x16mm (Covaris 520045)

Tubes are single use. They are used with rack 500111, Covaris E220 system. For the details of the Covaris system and tubes, see http://covarisinc.com/products/afa-ultrasonication/e-series/ and links from there.

- a computer with the following minimum requirements:

- 4 cores
- 16 Gb of memory
- 200 G of hard disk space
and the following tools installed:

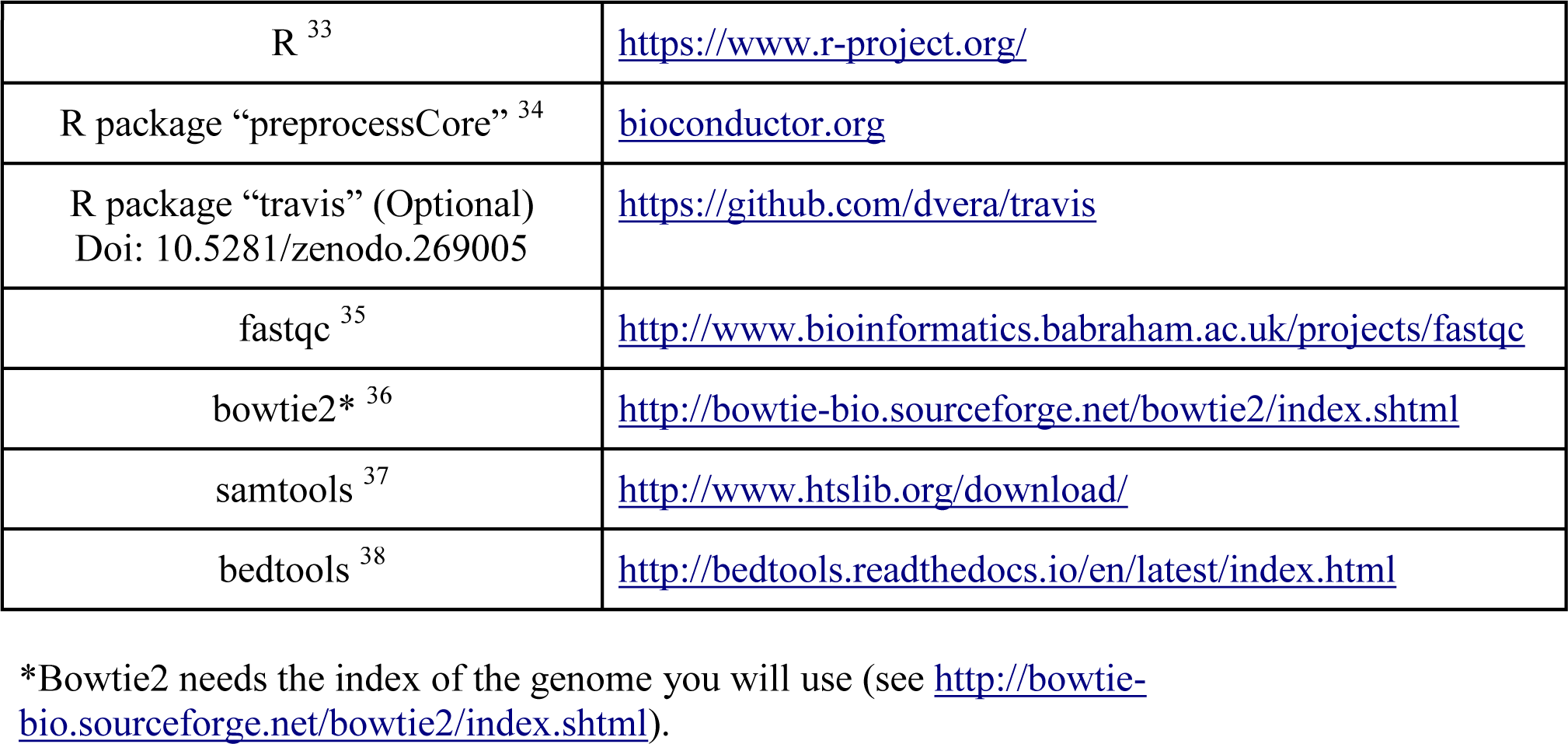

## PROCEDURE

### Step 1: BrdU pulse labeling and fixation of cells 3h

*This is for adherent cells in a T75 flask with 15 mL medium (for suspension culture in 15mL, skip steps 3 to 5).*

1. Add BrdU into medium to a final concentration of 100 μM
2. Incubate 2 hr at 37°C tissue culture incubator for BrdU incorporation
3. Rinse twice the cells gently with ice-cold PBS
4. Trypsinize cells with 2 mL of 0.2X Trypsin-EDTA, 2-3 min
5. Add 5 mL of FBS-containing medium, pipette gently but thoroughly, and transfer to 15 mL round bottom tube (Falcon 2059 or the like)
6. Centrifuge, 200 g, 5 min
7. Decant (or aspirate) supernatant carefully
8. Add 2.5 mL of ice-cold PBS / 1 *%* FBS, pipette gently but thoroughly CRITICAL STEP Double check the cell number at this point, after adding ethanol, it will be harder to count the cells since FBS makes precipitation.
9. Add 7.5 mL of ice-cold 100% EtOH, dropwise while gently vortexing (use the lowest rpm or hand shake the tube to avoid cell lysis by vigorous vortexing)
10. Seal cap, mix tube gently but thoroughly by inverting several times.
11. Store at -20°C until use (and in the dark since BrdU is light-sensitive). Lower temperature may cause freezing which damages cells. PAUSE POINT Fixed cells are stable in -20°C for more than a year if protected from light and evaporation.

Note: Starting from rapidly growing cells helps as they have a large percentage of S phase cells. In our experience, 2 million total cells, with >5% cells in S phase, yields enough early and late S phase cells for one replication assay (60,000 each) most of the time. Using round bottom tubes for cell fixation prevents cells from forming packed pellet that is hard to re-suspend later, but if small cell number is an issue, using conical tubes is fine.

### Step 2: FACS sample preparation and sorting 1.5h

*This is a "whole cell sorting" procedure. If the fixed cells’ condition is poor (making aggregates etc.) or cell sorter specification does not allow sorting whole cells (too narrow nozzle etc.), "nuclei preparation" procedure (see Supplementary method I.) may help.*

12. Transfer 2 × 10^6^ cells to a 15 mL conical tube.
13. Centrifuge at approximately 200 × g for 5 minutes at room temperature.
14. Decant supernatant carefully
15. Re-suspend the cell pellet in 2 mL 1% (vol/vol) FBS in PBS. Mix well by tapping the tube.
16. Centrifuge at approximately 200 × g for 5 minutes at room temperature.
17. Decant supernatant carefully.
18. Resuspend cell pellet in PBS / 1% FBS / PI / RNase A to reach 3 × 10^6^ cells/mL.
19. Tap the tube to mix and then incubate for 20 to 30 minutes at room temperature (22°C) in the dark. (count the cells during this time and adjust cell concentration if necessary)
20. Filter cells by pipetting them through 37-micron nylon mesh into a 5 mL polypropylene round bottom tube.
21. Keep samples on ice in the dark and proceed directly to FACS sorting. PAUSE POINT Alternatively, add 1/9 vol. DMSO and freeze in -80°C (light protected) until sorting. On sorting, thaw the cell suspension in a 37°C water bath. Removing DMSO is not necessary. Once thawed, keep the samples on ice in the dark.
22. Request FACS operator to collect 120,000 cells each of early and late S phase cells. (120,000 cells allow 6 reactions of BrdU IP). CRITICAL STEP It is hard to define the junction between G1 and S. Therefore, include some late G1 into the early S fraction. In addition, keep in mind that cells in late S phase proceed into G2 during the 2hr BrdU labeling time, hence, include early G2 in the late S fraction. Be sure to leave as small a gap as possible between the early and late S sorting windows. Otherwise, data for mid S replicating DNA will be missing and there is no reason to lose these cells. TROUBLESHOOTING

### Step 3: DNA preparation from FACS sorted cells 0.5h

23. Centrifuge the sorted cells at 400 × g or sorted nuclei at 800 × g for 10 minutes at 4°C.
24. Decant supernatant gently, only once (If the cell number is small, there may be no supernatant coming out by decanting, but do it for every sample for the consistency).
25. Add 1 mL of SDS-PK buffer containing 0.2 mg/mL Proteinase K every 100,000 cells collected and mix vigorously by tapping the tube. Seal around the tube cap with parafilm so that the caps would not pop out during the next step.
26. Incubate samples in a 56°C water bath for 2 hours.
27. Mix each sample thoroughly to get homogeneous solution after 56°C incubation, then aliquot 200 μl, equivalent to approximately 20,000 cells, into separate 1.5 mL tubes for each sample. (This one tube is for one library/IP. In order to see the consistency of IP, it is recommended to process at least 2 fractions per sample if any possible).
28. Add 800 μl Genomic Lysis Buffer from Zymo Quick-DNA Microprep kit and follow the kit protocol to purify DNA. Elute DNA into 50 μl H2O.

CRITICAL STEP Sometimes, you get more than 50 μl elution. This is because the wash buffer had not been completely removed from the column. Pay great attention to make sure you do not have wash buffer left on the column before adding 50 μl H2O for elution.

PAUSE POINT The purified DNA can be sheared immediately by Covaris or stored in -20°C.

### Step 4: Fragmentation 1h

29. The Covaris water bath needs to be chilled and de-gassed for 45 minutes before each use. See the manufacturer instructions for more information.
30. Using a 100-200 μl pipette tip, transfer the purified DNA from Step 28 into a microTUBE AFA Fiber (numbered on periphery of the cap) through the slit of the microTUBE. The slit closes automatically.
31. Keep the microTUBE AFA Fiber tube containing DNA on ice until fragmentation starts.
32. While de-gas is underway, set the shearing conditions. For 200 bp average fragment size, use 175W, 10% duty, 200 cycles/burst, 120 seconds, 7°C water.
33. Set the sample tubes on the rack and start shearing according to Covaris manual.
34. Once all tubes have been treated, spin the tubes at 600 RCF for 5 sec to collect all the liquid at the bottom of the tube.
35. Optional: Although Covaris is very reproducible, you may want to check the fragment size distribution, especially on your first try. In such case, concentrate the sheared DNA to 10-15 μL using DNA Clean & Concentrator-5 and check 1 μL on a Bioanalyzer HighSensitivity DNA chip.

### Step 5: Library construction 3h

*This basically follows NEB’s manual using 0.5 ng - 1 μg fragmented DNA. • Colored bullets indicate the cap color of the reagent to be added to a reaction.*

36. Mix the components in the table below in a 0.5ml PCR tube:

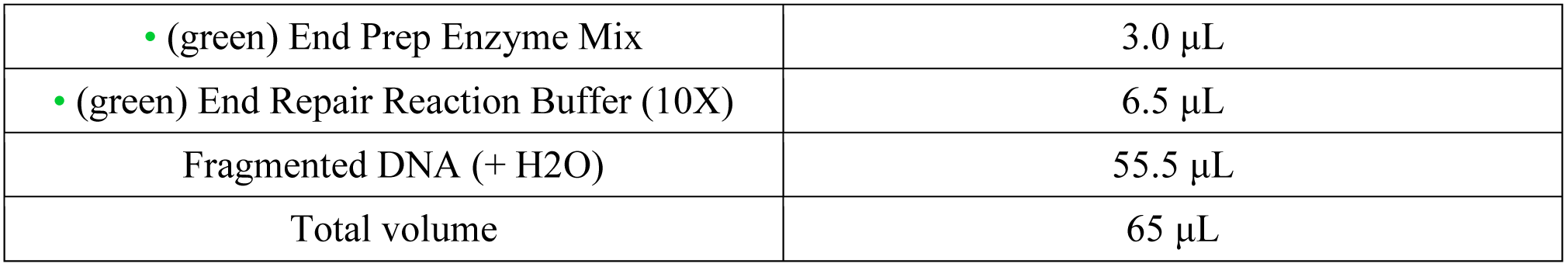
37. Mix by pipetting followed by a quick spin to collect all liquid from the sides of the tube.
38. Place in a thermocycler, after the lid is heated, and run the program of the table below.

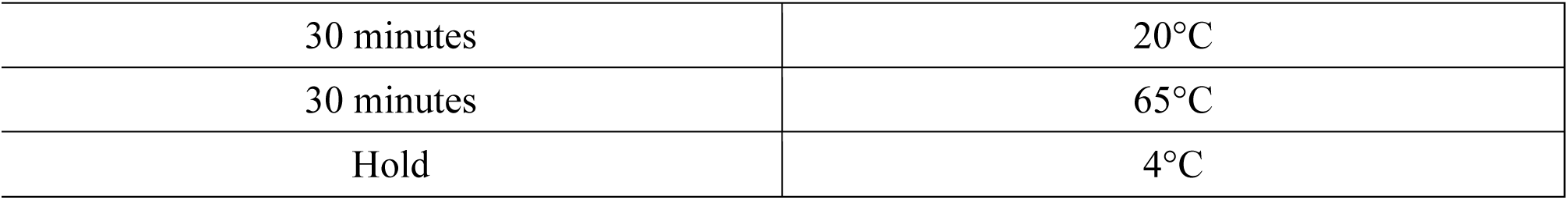
39. Add the components of the table below directly to the mix from Step 38 and mix well.

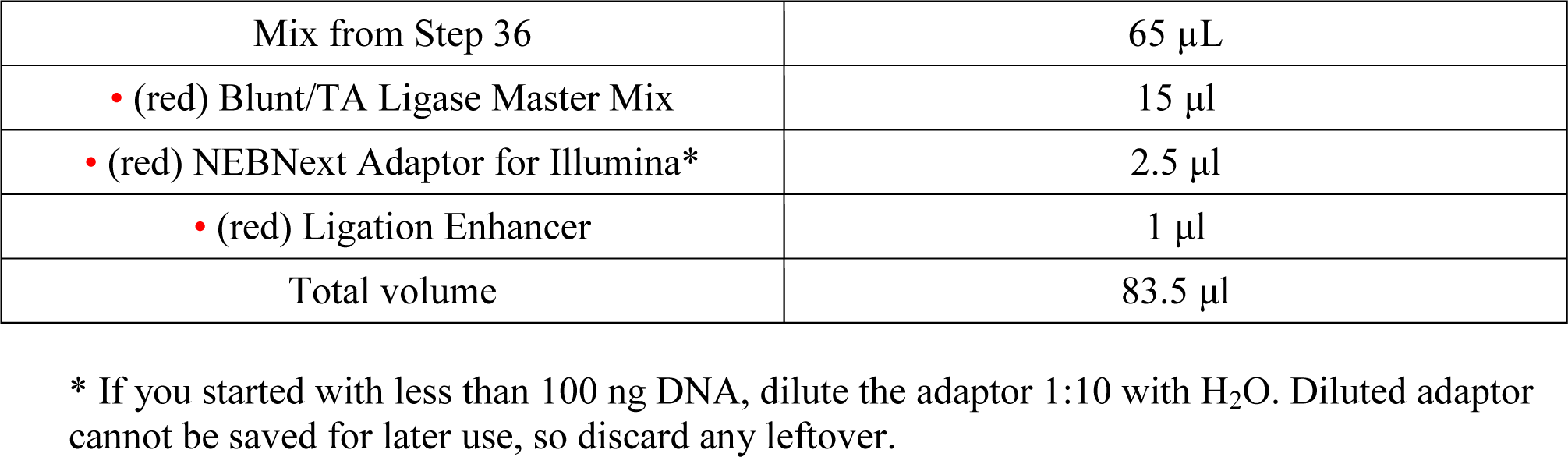
40. Mix by pipetting and then perform a quick spin to collect all liquid from the sides of the tube.
41. Incubate at 20°C for 15 minutes in a thermal cycler with heated lid off.
42. Add 3μl USER enzyme to the ligation mixture from Step 41.
43. Mix well and incubate at 37°C for 15 minutes with heated lid on.
44. Purify the DNA using DNA Clean & Concentrator-5. Elute into 50 μl H_2_O. PAUSE POINT The purified DNA can be stored at -20°C protected from light.

### Step 6: BrdU IP 1h and overnight

*The BrdU IP does not use beads to precipitate the antibodies / DNA-BrdU complexes. The centrifugation performed Step 54 is sufficient to pellet the complexes without need of beads because primary and secondary antibodies alone can make visible aggregates without BrdU-labeled DNA. However, visible precipitates do not guarantee successful capture of BrdU-labeled DNA.*

45. Add 450 μL TE to the DNA from Step 44.
46. Aliquot 60 μL 10X IP buffer to separate fresh tubes (one for each sample from Step 45).
47. Make 0.75 mL 1X IP buffer for each sample and start cooling on ice.
48. Denature DNA from Step 45 at 95°C for 5 minutes then cool on ice for 2 minutes.
49. Add the denatured DNA from Step 48 to the tube from Step 46.
50. Add 40 μL of 12.5 μg/mL anti-BrdU antibody.
51. Incubate 20 minutes at room temperature with constant rocking.
52. Add 20 μg of rabbit anti-mouse IgG. (Anti-mouse IgG concentration differs lot by lot. Check certificate of analysis.)
53. Incubate 20 minutes at room temperature with constant rocking.
54. Centrifuge at 16,000 × g for 5 minutes at 4°C
55. Remove supernatant completely (repeat pipetting and brief spin, first using 200 μL tips, finally using 10 μL tips).
56. Add 750 μ L of 1X IP Buffer that has been chilled on ice.
57. Centrifuge at 16,000 × g for 5 minutes at 4°C.
58. Remove supernatant completely, as in step 55.
59. Re-suspend the pellet in 200 μL digestion buffer with freshly added 0.25 mg/mL Proteinase K and incubate samples overnight at 37°C (air incubator).
60. Add 1.25 μL of 20 mg/ml Proteinase K to each tube.
61. Incubate samples for 60 minutes at 56°C (water bath).
62. Purify the DNA using DNA Clean & Concentrator -5 and elute into 16 μ L H_2_O.

### Step 7: Indexing and amplification 1.5h

63. Mix the components of the table below in 0.5mL PCR tubes . See supplemental data I for more information on NEBNext primers, and refer to NEB and Illumina manuals for the combination of index primers.

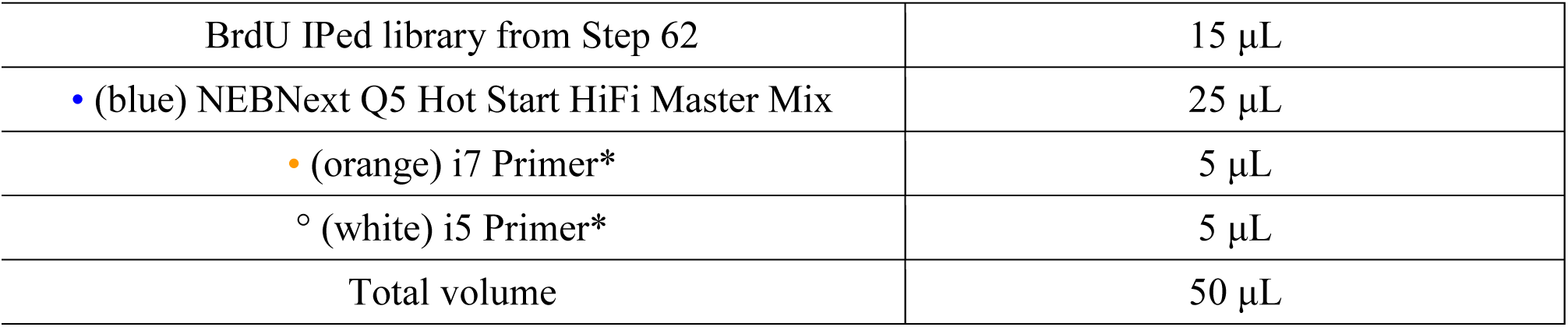 CRITICAL STEP Each library should get a unique combination of i7 and i5 primers.
64. Place in a thermocycler, after lid is heated, and run the program of the table below.

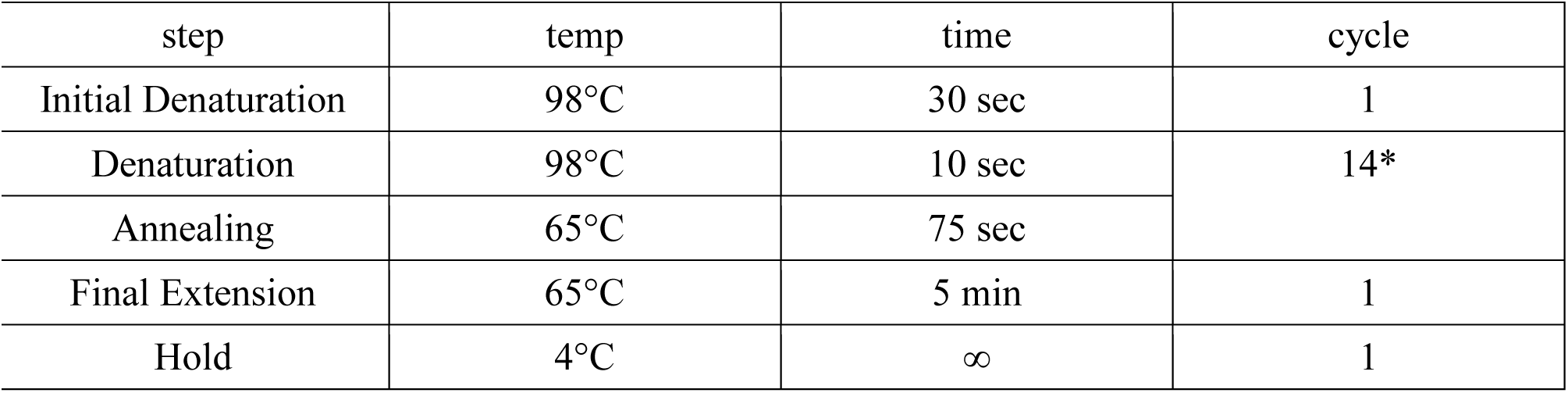
65. (Optional) In order to estimate the amount of your BrdU-IPed DNA and determine the optimal PCR cycle number for indexing, you can perform a qPCR using primers that anneal to adaptor region (in this case: NEBadqPCR_F; ACACTCTTTCCCTACACGACGC and NEBadqPCR_R; GACTGGAGTTCAGACGTGTGC) and serial dilution of your previous NGS library with known concentration as standard.
66. (Optional) You can run 5uL each of PCR reaction on a 1.5% agarose gel after PCR to check the size distribution by EtBr staining. A smear around 350 bp is expected. If no smear is detected, you need to re-amplify the reaction (the procedure for re-amplification is described in Step 78).

### Step 8: Purification 1h

67. Place AMPure XP beads at room temperature for at least 30 minutes.
68. Add H_2_O to each PCR reaction to make the final volume 100μL.
69. Vortex AMPure XP beads to resuspend.
70. Add 90μL of resuspended AMPure XP beads to the 100μ L PCR reaction. Mix well by pipetting up and down at least 10 times. CRITICAL STEP If you start from poorly fragmented DNA and you are sure you need to perform size selection at this moment, refer to NEB manual E7370 and optimize the volume of AMPure XP beads to use.
71. Incubate for 5 minutes at room temperature.
72. Quickly spin the tube and place the tube on an appropriate magnetic stand to separate the beads from the supernatant. Once the solution is clear (takes about 5 minutes), carefully remove the supernatant.
73. Mix ethanol and H_2_O to prepare fresh 80% ethanol (200 μL × [tube number + 1]). Add 200 μ L 80% freshly prepared ethanol to each tube while in the magnetic stand. Incubate at room temperature for 30 seconds, and then carefully remove and discard the supernatant.
74. Repeat Steps 72-73 two more times for a total of three washes.
75. Air dry the beads for 10 minutes while the tube is on the magnetic stand with the lid open but loosely covered by plastic wrap.
76. Elute the DNA from the beads
  (i) Add 33 μL 10 mM Tris-HCl. Mix well on a vortex mixer or by pipetting up and down.
  (ii) Quickly spin the tubes and return them to the magnetic stand.
  (iii) Once the solution is clear (takes about 5 minutes), transfer 31 μ L to a new tube. Store libraries at −20°C. PAUSE POINT Libraries can be stored indefinitely at -20°C

### Step 9: Quality control and pooling 5h

77. Check the DNA concentration using 1 μL of the library on Qubit dsDNA HS Assay Kit. (See Qubit manual). Expect 10-20 ng/μl. If the concentration is below detection, do another Qubit assay using 10 μL. This helps to determine the number of PCR cycles necessary for reamplification. If the DNA concentration is higher than 7 ng/μL, skip the Step 76.
78. If the DNA concentration is less than 7 ng/μl, you will likely have problems making the final 10 nM pool, so you will need to re-amplify the sample. Please note that it is better to use more PCR cycles from the beginning rather than re-amplifying, because re-amplification includes one more step of purification. Re-amplification can be done as following:

(i) Mix the components of the table below:

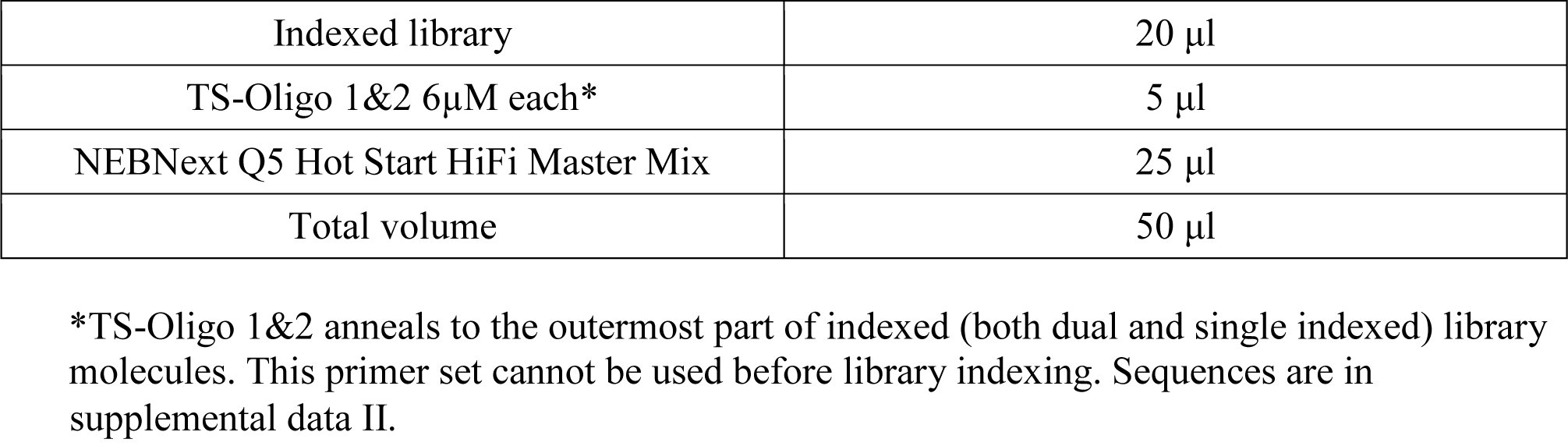
(ii) Place the mix in a thermocycler, after lid is heated, and run the program of the table below:

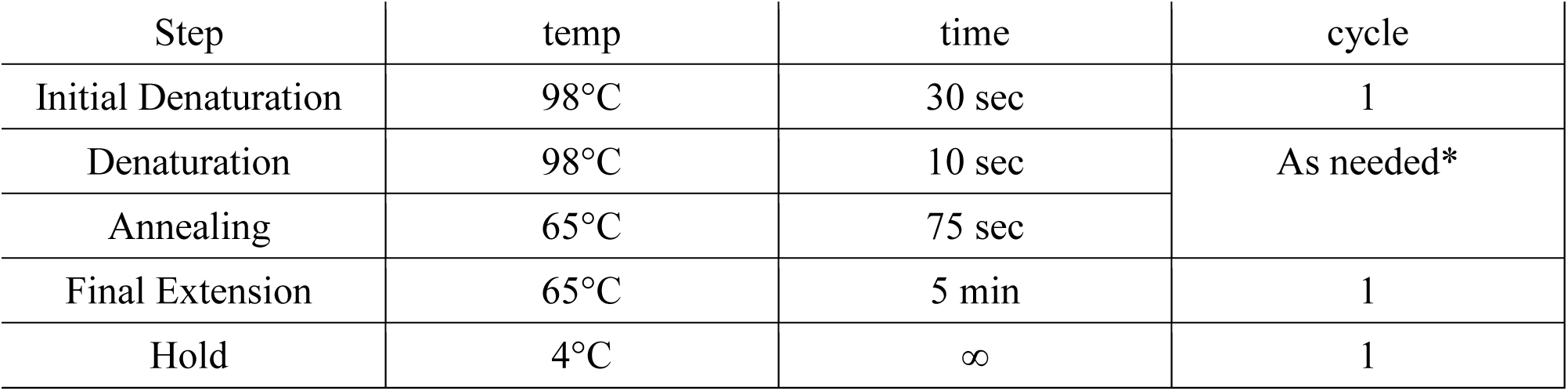 CRITICAL STEP Avoid over-amplification, since PCR cycles with low primer concentration causes library dimer, trimer, etc. formation.
(iii) Purify the PCR product as in Steps 67 to 76 and quantify DNA using Qubit. If DNA concentration is ≥ 10 ng/ul, proceed to Step 79. TROUBLESHOOTING
79. Using 2 ng/μ L dilutions of each library as templates, check the early and late fraction enrichment by PCR using primer sets listed by Ryba et al.18 . See supplemental data III for examples of Multiplex Primer mix.
  (i) Mix the components of the table below:

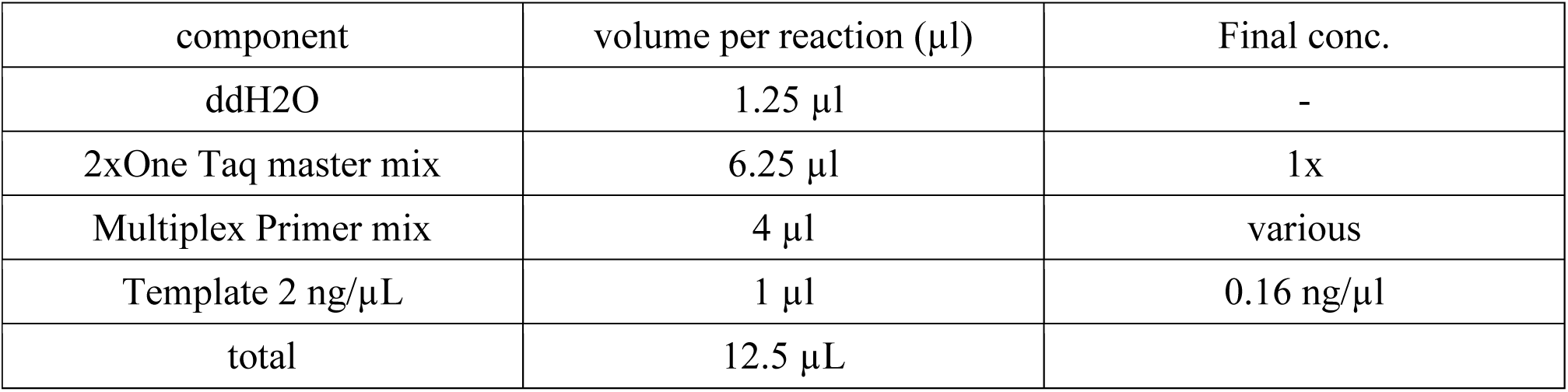
  (ii) Place the mix in a thermocycler, after lid is heated, and run the program of the table below:

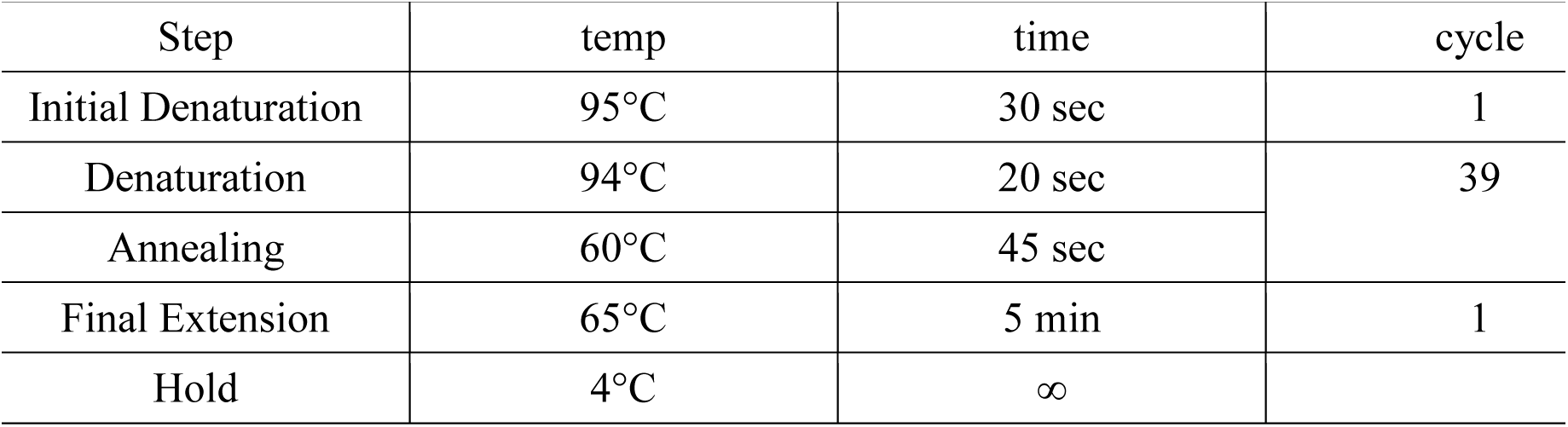 CRITICAL STEP Since the target size is small enough, this 2-step PCR works without an extension step. However, it is important to stick to this annealing condition (45s is relatively long) so that the primer can start extension slowly during the annealing instead of suddenly being stripped from the template by heating for the next cycle of denaturation.
  (iii) Run 4-5 μL of each PCR product (addition of loading buffer is not necessary for One Taq PCR) on 1.5% agarose gel containing EtBr to check the size of PCR products (specificity of PCR) and target enrichment. 30 minutes at 125V or 40 minutes at 100V is enough for this purpose (use the long gel, as you are separating multiplex PCR products). If the expected target enrichment is confirmed, proceed to Step 80. TROUBLESHOOTING
80. Check the size distribution of each library using the Bioanalyzer high sensitivity DNA kit or DNA 1000 kit according to your samples DNA concentration. See the Bioanalyzer/kit manual for details of the procedure, but first, place the reagents at room temperature. (~30 minutes for equilibration is recommended). See supplemental data IV for examples of good / bad quality libraries.
  (A) If your library concentration determined by Qubit is between 30 ng/μL and 50 ng/μL, load 1μL of each undiluted sample on a DNA 1000 chip. If the concentration is higher than 50 ng/μL (unlikely), dilute to 50 ng/μL and load 1μL on a DNA 1000 chip.
  (B) If your library concentration is less than 30 ng/μL, dilute to 0.5 ng/μL (based on Qubit) and load 1 μL to DNA High Sensitivity chip. So the aim is the upper limit of "quantitative range". The evaluation criteria for quality control results are:

(i) No adaptor/primer dimer peaks (below 150-180 bp) are detected. (Primer dimer contributes to clustering during sequencing but does not contribute to valid reads).
(ii) All the libraries to be pooled have similar and tight (~ 300 bp width) size distribution. (Molecules of different size have different clustering efficiency). If your samples do not meet these criteria, go back to Step 67. If all your samples pass these criteria, proceed to Step 81 (or Step 82 as Step 81 is optional).
81. (Optional) Determine of the molar concentration of each library by qPCR. This step may be omitted after you get used to the procedure. See manufacturer’s instruction for the latest update on KAPA qPCR kit. If your library has average 350 bp size and 10ng/uL concentration, the molar concentration is around 50nM, hence 1:10,000 dilution would fit within the standard curve. 10 μL reaction size works. Set up triplicates of standard DNA and duplicates of test samples (one dilution – usually 1:10,000, but decide the dilution factor based on your Qubit and Bioanalyzer results). After the PCR run, compensate and calculate the molar concentration of your samples as below using the average fragment size from Bioanalyzer.

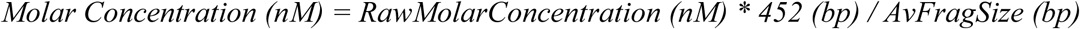

With *RawMolarConcentration,* the raw molar concentration from qPCR and *AvFragSize* your average fragment size (400 bp). *452* is the size of the standard DNA fragments used by the KAPA Illumina library quantification assay kit.
82. Pool the libraries. The number of libraries to pool is important to rich the good sequencing depth. For human or mouse samples, 5M mapped reads per library give usable data, which correspond to less than 10M sequenced reads (depending on the quality of the sequencing). Our HiSeq2500 generates ~160M reads per lane, so we usually pool 12 to 16 libraries per lane. Also, pooling less than 4 indexes is not recommended due to the low complexity of the index.
  (i) Using 0.05 % Tween 20 in 10 mM Tris pH 8, adjust each library to 10 nM (if you skipped the Step 81, estimate the molar concentration of each library using Bioanalyzer region function).
  (ii) Mix equal volume of each 10 nM library to make a pool. One pool fills one lane. CRITICAL STEP Make at least 23 μL of each pool (preferably ~50 μL for backup) in a low-adhesion tube so that DNA concentration would not easily change by adsorption while waiting for the run.
  (iii) (Optional) Add 1% PhiX spike into the pool (If you submit the pool to sequencing core etc. for service, PhiX is most likely to be added at the sequencing core) . PhiX library is made by Illumina and works as a positive control of the run itself (clustering and sequencing reactions). PAUSE POINT The pool can be stored in -20°C.
83. Quality control of the pool(s): take an aliquot from each pool and adjust them to 0.5 ng/μL (based on Qubit quantification), and run 1 μL of each on a Bioanalyzer DNA HiSensitivity chip to determine the average fragment size of the pool
84. Sequence the pooled libraries on a Hi-seq Illumina sequencer. Generally, 50bp single end reads is sufficient for normal samples covering the unique sequences of the genome. However, longer reads may be required to parse alleles by SNPs.

### Step 10: Data Analysis 1 day

*Data analysis is performed on fastq or fastq.gz files, and lead to the production of genomic coverage files with the log ratio of early versus late samples.*

CRITICAL STEP This pipeline is written to process files with names that follow a specific nomenclature: early and late sequencing data originated from the same sample must have matching name, with an "_E_" before the extension of the file from early S cells, and an "_L_" before the extension of the file from late S cells. If you use paired-end sequencing data, pairs files are here named with R1 and R2 before the extension. An example of a single-end fastq file name could be “my_sample_1_E_.fastq”, and a pair of paired-end fastq files name could be “my_sample_1_E_R1.fastq” and “my_sample_1_E_R2.fastq”.

85. Control the quality of the reads using fastqc. To install fastqc, see http://www.bioinformatics.babraham.ac.uk/proiects/fastqc for the installation and use of this tool. TROUBLESHOOTING
86. Process the fastq files. If you have multiple fastq files per libraries, see supplemental method II to catenate the fastq files. See supplemental method III to generate log ratio coverage files using R, and supplemental method IV to map the reads on two genomes and generate the log ratio coverage files for each genome using R. This step require chrom.size file of the genome you will you use for the analysis. Many of these files can be directly downloaded from UCSC server at ftp://hgdownload.cse.ucsc.edu/goldenPath/. This file contains two columns: each chromosome name and size in bp, separated by a tabulation. During this step, you can specify the size of the genomic windows you will use, depending on the further analysis you will perform on the RT datasets. We use genomic windows from 5 to 50kb. You can also specify the step between windows, to generate overlapping (or not) windows for example. CRITICAL STEP This file must be sorted on the first column, following alphabetic order (for example, “chr10” will be before “chr2”).
  (i) Create the following script.sh text file in a text editor, and register it in the fastq files directory
    (A) If you are analysing single-end data

~~~
*# Generating 50kb windows positions along the genome. You can change the -w value to change the windows size, and the -s value, to change the steps (to make overlapping windows for example)*
*sort -k1,1 -k2,2n* ***your_genome.chrom.sizes*** *>* ***your_genome_sorted.chrom.sizes*** *bedtools makewindows -w 50000 -s 50000 -g* ***your_genome_sorted.chrom.sizes*** *>* ***your_genome_windows.bed***

*# Mapping fastq(.gz) and making bed with values on genomic windows for file in *.fastq*; do*
*# Mapping*
*(bowtie2 -x* ***/path/to/your/genome*** *--no-mixed --no-discordant --reorder -U $file -S ${file%.fastq*}.sam 2>> mapping_log.txt*
*# sam to bam conversion*
*samtools view -bSq 10 ${file%fastq*}.sam > ${file%.fastq*}.bam*
*# bam to bed conversion*
*bamToBed -i ${file%.fastq*}.bam | cut -f 1,2,3,4,5,6 | sort -T. -k1,1 -k2,2n -S 5G > ${file%.fastq*}.bed*
*# bed line number calcul*
*x=*’*wc -l${file%.fastq*}.bed* | *cut -d′′-f l*’
*# generate coverage on genomic windows*
*bedtools intersect -sorted-c -b ${file%.fastq*}.bed -a* ***your_genome_windows.bed*** | *awk -vx=$x ′{print $1,$2,$3,$4*1e+06/x}′ OFS=′\t′ > ${file%.fastq*}.bg) & done*
*wait*

*# Calculating RT*
*for file in *_E_.bg; do*
*paste $file ${file%E_.bg}L_.bg* | *awk ′{if($8 != 0 && $4 != 0){print $1,$2,$3,log($4/$8)/log(2)}}′ OFS=′\t′ > ${file%E_.bg}T_.bg done*

*# Merging RT files*
*bedtools unionbedg -filler "NA" -i *T_.bg > merge_RT.txt*
~~~
    (B) If you are analysing paired-end data

~~~
*# Generating the 50kb windows positions along the genome. You can change the -w value to change the windows size, and the -s value, to change the steps (to make overlapping windows for example)*
*sort -k1,1 -k2,2n* ***your_genome.chrom.sizes*** *>* ***your_genome_sorted.chrom.sizes*** *bedtools makewindows -w 50000 -s 50000 -g* ***your_genome_sorted.chrom.sizes*** *>* ***your_genome_windows. bed***

*# Mapping fastq(.gz) and making bed with values on genomic windows for file in *R1.fastq*; do*
     *# Mapping*
     *(bowtie2 -x* ***/path/to/your/genome*** *--no-mixed --no-discordant --reorder -1 $file -2 ${file%R1.fastq*}R2.fastq* -S ${file%R1.fastq*}.sam 2>> mapping_log.txt*
     *# sam to bam conversion*
     *samtools view -bSq 10 ${file%R1.fastq*}.sam > ${file%R1.fastq*}.bam*
     *# bam to bed conversion*
     *bamToBed -i ${file%R1.fastq*}.bam* | *cut -f 1,2,3,4,5,6* | *sort -T. -k1,1 -k2,2n -S 5G > ${file%R1.fastq*}.bed*
     *# bed line number calcul*
     *x= ‘wc -l ${file%R1.fastq*}.bed* | *cut -d′* ′ *-f 1’*
     *# generate coverage on genomic windows bedtools intersect -sorted-c -b ${file%R1.fastq*}.bed-a* ***your_genome_windows.bed*** | *awk - vx=$x ′{print $1,$2,$3,$4*1e+06/x}′ OFS=′\t′ > ${file%R1.fastq*}.bg) &*
*done*
*wait*
*# Calculating RT*
*for file in *_E_.bg; do*
*paste $file ${file%E_.bg}L_.bg* | *awk ′{if($8 != 0 && $4 != 0){print $1,$2,$3,log($4/$8)/log(2)}}′ OFS=′\t′> ${file%oE_.bg}T_.bg done*
*# Merging RT files*
*bedtools unionbedg -filler "NA" -i *T_.bg > merge_RT.txt*
~~~ The bold paths and names have to be adapted to the path and names used in your computer. For more information on the genome path used by bowtie2 and others bowtie2 options, see http://bowtie-bio.sourceforge.net/bowtie2/manual.shtml. For better performance, you can allow bowtie2 multiple processors, depending on your resources, with the option -p [number of processors] (see bowtie2 documentation). If reads quality is bad at the end of the reads, you can trim the reads with bowtie2 option --trim3.
  (ii) Open a shell and go to the fastq files directory:

~~~
  $ cd path/to/files/
~~~
  (iii) Make the script.sh file executable:

~~~
  $ chmod 755 script.sh
~~~
  (iv) Execute the script.sh file:

~~~
  $ ./script.sh
~~~
87. Post-process the bedgraph files. This step is performed in R, using the package “preprocessCore”
  (i) Open R and go to the bedgraph files directory:

~~~
  > setwd("path/to/files/")
~~~
  (ii) Load the “preprocessCore” package:

~~~
  > library(preprocessCore)
~~~
  (iii) Import the merged bedgraph files:

~~~
  > merge<-read.table("merge_RT.txt", header=FALSE)
  >colnames(merge)<-c(c("chr","start","end"),list.files(path=".",pattern="*T_.bg"))
  > merge_values<-as.matrix(merge[,4:ncol(merge)])
~~~
  (iv) Set the datasets to use for quantile normalization (bold names have to be adapted):

(A) normalization on all datasets:

~~~
     > ad<-stack(merge[,4:ncol(merge)])$values
~~~
(B) normalisation on one datasets:

~~~
     > ad<-merge[,"**my_sample_T_.bg**"]
~~~
(C) normalisation on multiple datasets (You can add as many datasets as you want):

~~~
     >ad<-stack(merge[,c("**my_sample_1_T_.bg**","**my_sample_2_T_.bg**")])$values
~~~
  (v) Normalise the data:

~~~
   > norm_data<-normalize.quantiles.use.target(merge_values,ad)
   > merge_norm<-data.frame(merge[,1:3],norm_data)
   > colnames(merge_norm)<-colnames(merge)
~~~
  (vi) Register the quantile normalized data into bedgraph files:

~~~
   > for(i in 4:ncol(merge_norm)){write.table(merge_norm[complete.cases(merge_norm[,i]), c(1,2,3,i)], gsub(".bg", "qnorm.bedGraph", colnames(merge_norm)[i]), sep="\t",row.names=FALSE, quote=FALSE, col.names=FALSE)}
~~~
  (vii) Select the chromosome for Loess smoothing (You can modify the pattern option to select different chromosomes. Here, the pattern selects all chromosomes except chromosomes containing “_” in their name (which can be problematics for Loess smoothing), and the chromosomes Y and M (mitochondrial)):

~~~
  > chrs=grep(levels(merge_norm$chr),pattern=" [_YM] ",invert=TRUE,value=TRUE)
~~~
  (viii) (Optional) Check the list of selected chromosomes:

~~~
  > chrs
~~~
  (ix) Initialise an R-list to stock your datasets:

~~~
  > AllLoess=list()
~~~
  (x) Perform Loess smoothing (This smoothing is similar to the Loess smoothing used by Ryba et al 18. for repli-chip analysis). The window size used for span value (in bold) can be adapted. We are using here 300kb windows but you can increase this value to increase the smoothing.

~~~
  > for(i in 1:(ncol(merge_norm)-3)) {
AllLoess[[i]]=data.frame();
cat("Current dataset:", colnames(merge_norm)[i+3], "\n");
for(Chr in chrs){
RTb=subset(merge_norm, merge_norm$chr==Chr); lspan=**300000**/(max(RTb$start)-min(RTb$start)); cat("Current chrom:", Chr, "\n");
RTla=loess(RTb[,i+3] ~ RTb$start, span=lspan);
RTl=data.frame(c(rep(Chr,times=RTla$n)), RTla$x, merge_norm[which(
merge_norm$chr==Chr & merge_norm$start %in% RTla$x),3],RTla$fitted); colnames(RTl)=c("chr","start","end",colnames(RTb)[i+3]); if(length(AllLoess[[i]])!=0){
AllLoess[[i]]=rbind(AllLoess[[i]],RTl)};
if(length(AllLoess[[i]])==0){
AllLoess[[i]]=RTl}}}
~~~
  (xi) Register the Loess smoothed data into bedgraph files:

~~~
  > for(i in 1:length(AllLoess)){write.table(AllLoess[[i]][complete.cases(AllLoess[[i]]),],
gsub(".bg","Loess.bedGraph", colnames(AllLoess[[i]]))[4], sep="\t", row.names=FALSE, quote=FALSE, col.names=FALSE)}
  You can now exit R.
~~~
88. (Optional) Merge your bedgraph files for further analysis.
  (i) Open a terminal and go to your bedgraph repertory:

~~~
  $ cd path/to/files/
~~~
  (ii) Merge the loess smoothed bedgraph files:

~~~
  $ bedtools unionbedg -filler "NA" -i *Loess.bedGraph > merge_Loess_norm_RT.txt
~~~
89. Your RT data are now registered into bedgraph files into your repertory. You can visualize them using a genome viewer (IGV, IGB, UCSC genome browser) or on our replication domain platform (http://www.replicationdomain.org). You can also perform further analysis, as identifying the early and late domains and comparing RT between your samples, as described by Ryba et al. 18.

TIMING

**Step 1: BrdU pulse labeling and fixation of cells** 3h

**Step 2: FACS sample preparation and sorting** 1.5h

**Step 3: DNA preparation from FACS sorted cells** 0.5h

**Step 4: Fragmentation** 1h

**Step 5: Library construction** 3h

**Step 6: BrdU IP** 1h and overnight

**Step 7: Indexing and amplification** 1.5h

**Step 8: Purification** 1h

**Step 9: Quality control and pooling** 5h

**Step 10: Data Analysis** 1 day

## TROUBLESHOOTING

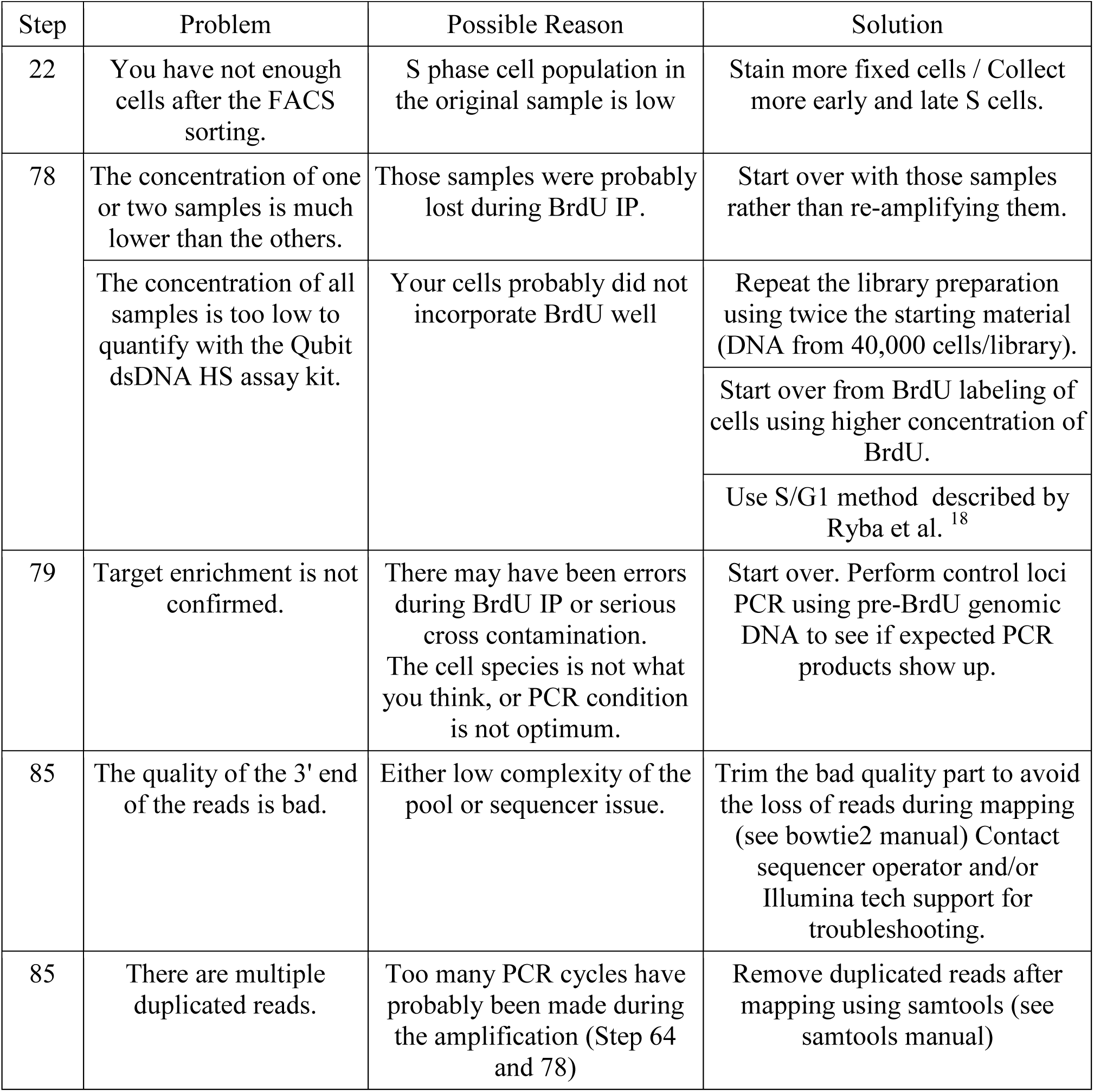

## ANTICIPATED RESULTS

### Reproducibility

DNA RT is a robust epigenetic property of specific cell types. Perturbations of the cells by knockout or knock-down experiments have a relatively weak impact on this program ^2,39,40,41,42^ Thus, assessing how a condition can affect this program can be challenging. Our method is highly reproducible, producing high correlations between replicates to the extent that small differences between samples can be detected and, if the intent is to compare a lot of different experimental conditions, one can be quite confident with a single experiment, providing the quality control standards are met. Our successive steps of normalization make comparisons of even closely related specimens quite facile (Figure 2), allowing high confidence identification of differences between samples.

**Figure 2:**
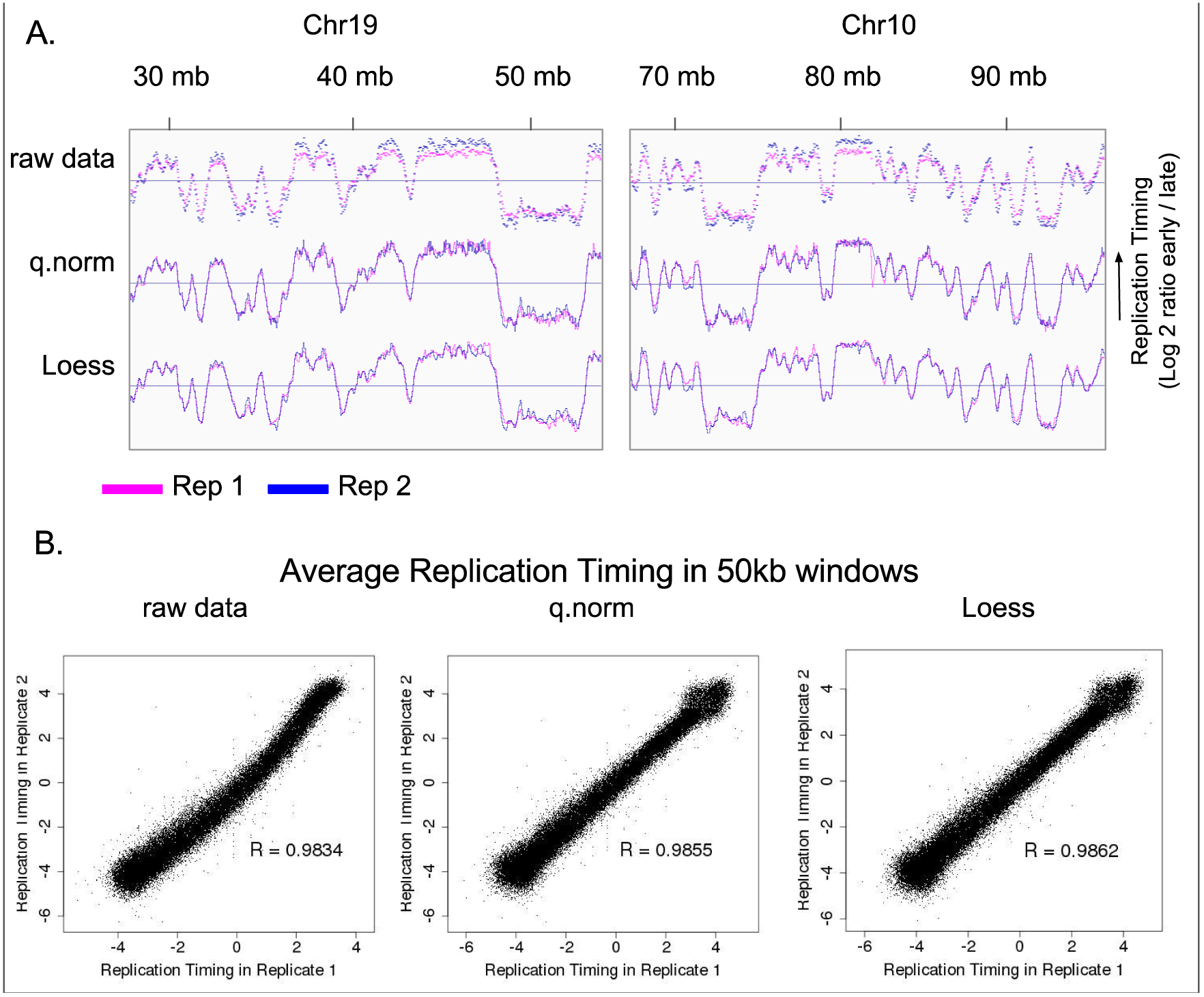
Quantile normalization and Loess smoothing allow comparison between samples. A.: Before and after each normalization steps Replication timing (RT) profile of two Repli-seq replicates (F121-9 mouse ESC, mapped on mm10). Data are visualized using IGV. B.: Correlation between 50kb windows along the genome of samples in A. R = Pearson correlation coefficient, q.norm = quantile normalization, Loess = Loess smoothing.

### Accuracy

Repli-chip is a well-accepted method to study RT. We have shown that repli-seq gives similar results to repli-chip, in both human and mouse cells 15 (Figure 3), and that comparisons of genomewide RT can be made between repli-seq and repli-chip. The two platforms can even be combined for clustering experiments. The advantages of repli-seq are the genomic coverage (which includes repetitive sequences should those be desirable to analyze), which is dependent only upon the mappability and sequencing depth, the ability to distinguish SNPs or parse phased genomes, and the ability to analyze any species with the same method.

**Figure 3.**
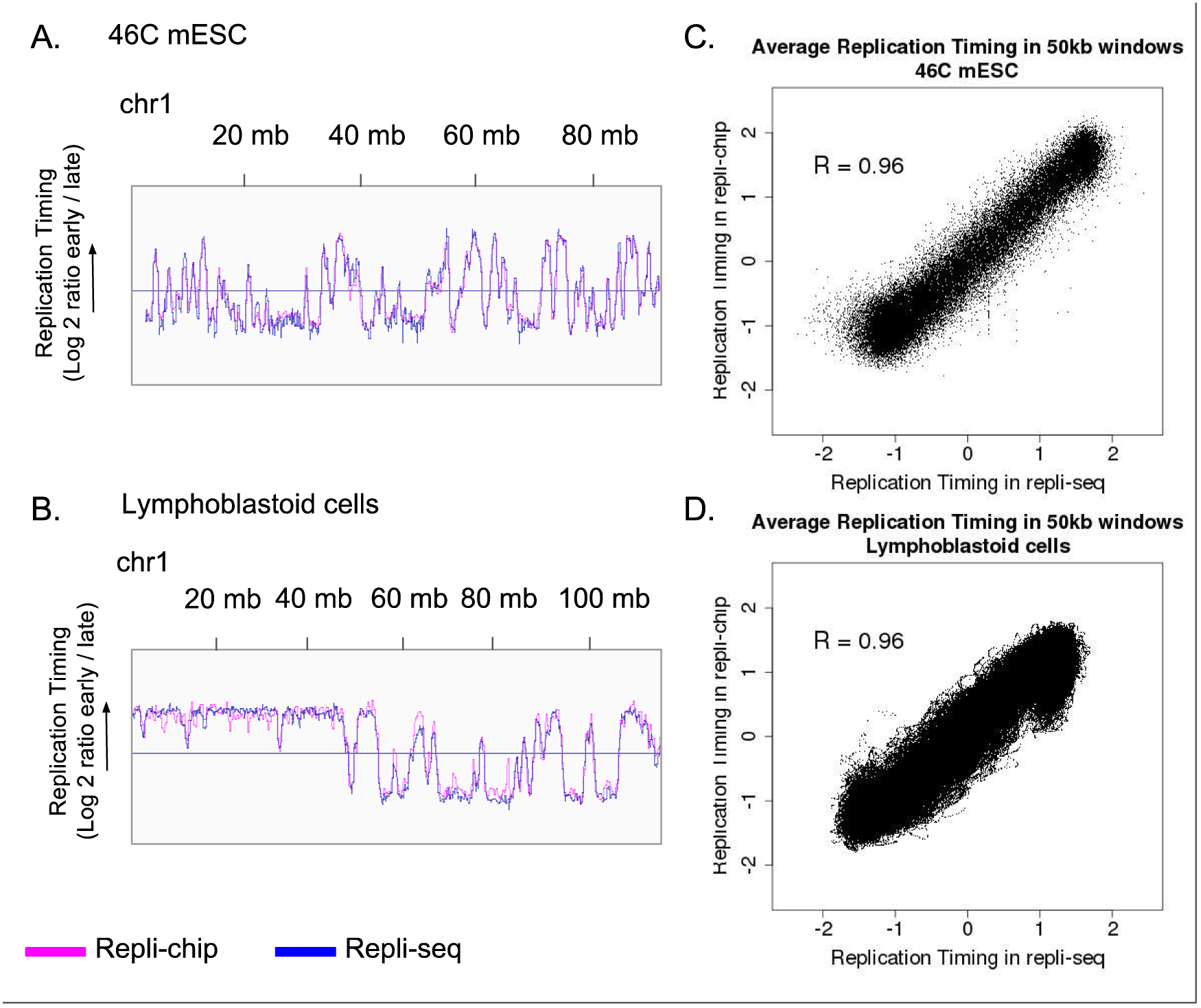
Repli-chip and Repli-seq give highly similar replication timing profiles at genome-wide level. A. B.: Replication timing (RT) profile on a portion of chrl of 46C mouse ESC (mm10) (A.) and human lymphoblastoid cells (hg38) (B.), visualized using IGV. RT is defined as the log2 ratio early fraction on late fraction (reads number is normalized on number of mapped reads for repli-seq). C. D.: correlation between average replication timing on 50kb windows on the whole genome in 46C mouse ESC (C.) and human lymphoblastoid cells (D.). Data are scaled in R prior to visualisation. R = Pearson correlation coefficient.

### Sequence specificity

The sequence information obtained from repli-seq can be used to discriminate between homologous regions, if the sequencing breadth is appropriate for the polymorphism density (e.g. parsing genomes with a SNP density of 1/1000 bp such as normal human variation requires 10 times the sequencing reads), permitting the study of differences in RT between homologous regions 21. We show here an example using hybrid mouse embryonic stem cells (Figure 4) with SNP density of ~1/100 bp. Discrimination in these cells of the original strain of homologous regions reveals subtle differences between homologous regions. Such regions could be compared with transcription data to study the link between transcription and RT. Repli-seq could also be used to discriminate between the RT of two X chromosomes in hybrid female cells, and the impact of X chromosome activation status on RT.

**Figure 4.**
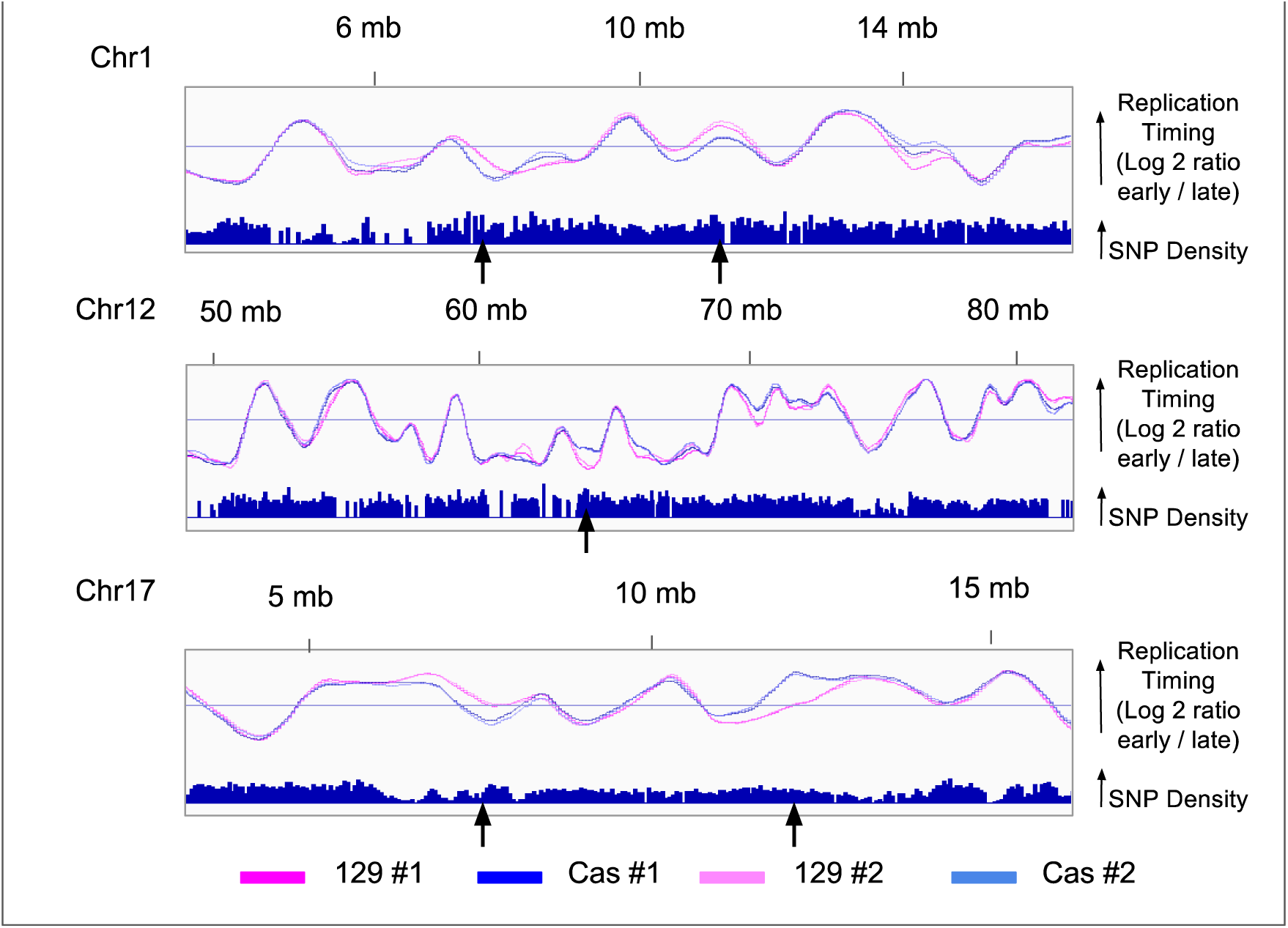
Repli-seq allows the discrimination between haplotypes. Comparison of replication timings of three homologous regions in mouse hybrid cells 129-cas. Reads have been mapped to cas and 129 reference genomes respectively. Arrows show major changes. Data are visualized using IGV.

## DATA ACCESSION

Data used to generate the figures are available on GEO (http://www.ncbi.nlm.nih.gov/geo/), under the numbers GSE37987 and GSE95092.

## AUTHORS CONTRIBUTIONS STATEMENTS

D.G., C. M. and T.S. conceived the study and designed the experiments.

T.S., K.W., J.S., C.T.G., C.N., E.N. and J.C.R.M. performed wet experiments.

D. V., J. S. and C.M. devised the computational methods.

C.M., T.S., and D.G. wrote the manuscript.

## ACKNOWLEDGMENTS

We thank Ruth Didier for assistance in cell sorting. This work was supported by NIH GM083337, GM085354, DK107965 to DMG. CM is supported by ARC French fellowship SAE20160604436.

## COMPETING FINANCIAL INTERESTS

The authors declare that they have no competing financial interests.

